# Extending the least-cost theory of stomatal regulation to include soil moisture stress

**DOI:** 10.64898/2026.06.23.733952

**Authors:** David Sandoval, Victor Flo, Han Zhang, Iain Colin Prentice

## Abstract

- Terrestrial biosphere models commonly use empirical scaling factors to represent soil moisture constraints on carbon and water fluxes, but these lack mechanistic grounding and produce inconsistent estimates of soil moisture limitations on primary production and transpiration across models.
- Here we extended the least-cost hypothesis for optimal stomatal conductance to account for soil moisture limitations by allowing soil water availability to modulate the carbon cost of water transport, drawing on the observed temperature dependence of stem respiration, and derived a simple empirical approximation to the theory using global δ^13^C and eddy covariance data.
- The empirical analysis shows moderated thermal acclimation of stem respiration and a weak increase in water transport costs with aridity, supporting the interpretation that the decline in light-use efficiency (LUE) under arid conditions is primarily attributable to non-stomatal limitations.
- Validation against an independent global dataset of sapflow-derived canopy conductance and transpiration shows that the revised scheme significantly improves the predictive power of the least-cost hypothesis, offering a more mechanistically coherent alternative to existing soil moisture parameterisations.

## Introduction

Explaining how soil moisture limitation constrains carbon uptake and transpiration remains a major challenge for terrestrial biosphere models (TBMs) partly because drought responses emerge from coupled changes in stomatal regulation, plant hydraulics and photosynthetic metabolism (Harrison et al., 2021; Prentice et al., 2015; Rogers et al., 2017). The basic mechanisms underlying the progression of this limitation are understood, at least qualitatively. A reduction in soil water potential (Ψ_s_) triggers accumulation of abscisic acid in the leaf apoplast, causing stomatal closure and immediately reducing both assimilation and transpiration (Buckley, 2019; Farquhar and Sharkey, 1982). Reduced transpiration increases leaf temperature and hence the leaf-to-air vapour pressure deficit (*D*), increasing the demand for water uptake. If water demand exceeds supply, the impairment of transpiration and water uptake causes dehydration (Zhang and Bartels, 2018) and a reduction in the relative leaf water content (RWC). With less water, the concentration of solutes increases, reducing the leaf water potential Ψ_l_ (Bartlett et al., 2012; Schulze et al., 2019). Ψ_l_ also decreases with increasing leaf temperature (Nobel, 1983). If there is sufficient water in the soil, this reduction in Ψ_l_ increases the gradient (ΔΨ) causing a negative feedback with water uptake, moving the system back to a steady state. However, if there is not sufficient water in the soil, the loss of soil conductivity will greatly reduce the uptake and the system will not return to a steady state; further dehydration then risks causing embolism and reduction in the plant-level hydraulic conductance (*K*_p_) (Lens et al., 2022; Sanchez-Martinez et al., 2020).

Generally, under moderate dehydration, when leaf relative water content remains above approximately 70–80%, reductions in assimilation are mainly attributable to stomatal closure and the resulting decline in intercellular CO_2_ concentration (*c*_i_) (Flexas, 2002), while maximum carboxylation capacity and quantum yield often remain largely reversible upon rehydration (Bhattacharya, 2021; Martín-Gómez et al., 2023; Zhang and Bartels, 2018). Under more severe or prolonged dehydration, however, non-stomatal limitations become increasingly important. These limitations include mesophyll conductance and metabolic processes such as electron transport, RuBP regeneration, Rubisco activity and photochemical efficiency. The excess in excitation energy caused by this metabolic impairment promote reactive oxygen species formation and membrane damage (Hu et al., 2023; Meyer and Genty, 1999; Murata et al., 2007; Zhang and Bartels, 2018). These metabolic and structural effects can limit post-drought recovery and therefore cannot be captured by stomatal limitation alone.

Plants have developed multiple adaptations to deal with soil moisture stress, depending on its severity and duration (Volaire, 2018). In the long term, arid climates favour plants with sclerophyllous leaves, with higher elastic moduli that can produce larger ΔΨ and extract more water from dry soils (Harrison et al., 2010). Other plants in arid climates have developed vascular systems with increased tolerance for low water potentials, reducing the risk of embolism (Franklin et al., 2023; Olson et al., 2018). In climates subject to seasonal transitions between dry and wet seasons, deciduousness is a common “avoidance” strategy to reduce dry-season transpiration (Kunert et al., 2021). Shifting carbon allocation to root biomass is a common “tolerance” strategy allowing survival through the dry season (Bachofen et al., 2024a). These strategies imply that drought responses are not cost-free: maintaining hydraulic safety, water uptake capacity and drought tolerance requires carbon investment. On shorter timescales, plants rely on a coordination between physiology and metabolism. The least-cost hypothesis, as defined by Prentice et al. (2014) , provides a useful theoretical framework for this coordination. According to the least-cost hypothesis, this coordination sets the ratio of leaf-internal to ambient CO_2_ (χ = *c*_i_/*c*_a_) in such a way as to minimize the combined costs of maintaining the capacities for carbon assimilation and associated water transport (Harrison et al., 2021; Prentice et al., 2014). As currently formulated, however, this hypothesis predicts only the reduction of χ with increasing atmospheric dryness (vapour pressure deficit) and does not consider how this response is modified by drought.

To include soil moisture (θ) constraints on stomatal conductance, some current models use plant hydraulic theory and water potentials to formulate resistances to water transport (Paschalis et al., 2024) or as a criterion to find the optimum stomatal opening (Sabot et al., 2022; Wang et al., 2020c). Hydraulic theory for trees is well established, and xylem hydraulic conductances and vulnerability curves have been measured for many species. Scaling up from individual trees to ecosystems for global modelling purposes is challenging, however. Most models designed for global application treat soil moisture stress more simply, by applying a multiplicative factor β(θ) ∈ [0,1] that penalises fluxes and/or conductance(s) as a function of relative soil moisture or relative drop in water potential (Medlyn et al., 2016; Rogers et al., 2017; Seneviratne et al., 2010a). Although the β(θ) method provides a pragmatic way to implement soil moisture stress, it suffers from three problems.

First, different models define the relative soil water availability drop (θ − θ_*low*_)⁄(θ_*upp*_ − θ_*low*_) in different ways. For example, the upper limit (θ_upp_) may be set at the saturation value of θ for a given soil, as in JSBACH (Reick et al., 2021); or at the value corresponding to a water potential of –33 kPa, as in JULES (Harper et al., 2021); or occasionally at other values, such as –10 kPa for sandy soils in HTESSEL (ECMWF, 2019). The lower limit (θ_low_) is typically assumed to be the value of θ at –1.5 MPa (Seneviratne et al., 2010b), taken to be the wilting point – although in reality this point varies among species. Inconsistent thresholds are also applied if Ψ_s_ is used as the control variable instead of θ (Rogers et al., 2017). β(θ) can adopt different shapes by incorporating an exponent, resulting in concave or convex curves (Metselaar and de Jong van Lier, 2007; Verhoef and Egea, 2014). It can be applied to different quantities – directly to gross primary production (GPP), or to transpiration, stomatal conductance or maximum carboxylation capacity (*V*_cmax_). Sometimes multiple β(θ) are applied to different quantities (De Kauwe et al., 2013) and in some cases, the parameters of these curves are prescribed separately by plant functional types (PFTs), but there is little empirical support for the imposed differences between PFTs (Medlyn et al., 2016).

Second, applying an identical β(θ) factor to both GPP and *g*_s_ fails to reproduce observed effects of soil moisture on water use efficiency (WUE; the ratio of assimilation to transpiration) (Niu et al., 2011; Poppe Terán et al., 2023) or intrinsic water use efficiency (iWUE, the ratio of assimilation to stomatal conductance) (Jiang et al., 2024a; Li et al., 2022a), because β(θ) cancels out algebraically. In other words, a multiplicative stress factor may reduce the magnitude of fluxes, but it does not necessarily alter their ratio. This is illustrated by equation (1), based on the least-cost formalism:

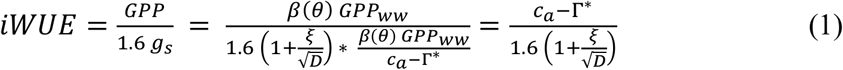

where GPP_ww_ is the ‘well-watered’ GPP, ξ is the control parameter in the least-cost model, *c*_*a*_ is the atmospheric CO_2_ concentration, and Γ* is the photorespiratory compensation point. Moreover, applying a β(θ) factor to GPP or *g*_s_ does not allow θ to modify the sensitivity of g_s_ to *D*, as observed experimentally (Asargew et al., 2024), and inferred using remote sensing in the Amazon (Green et al., 2020; Zhang et al., 2023).

Third, however it is applied, the β(θ) approach cannot predict metabolic acclimation to low soil moisture – a process by which *V*_cmax_ increases in response to lowered *c*_i_ (Mengoli et al., 2022; Ren et al., 2024). This phenomenon can be understood as a consequence of the coordination hypothesis, which states that *V*_cmax_ is set at the level permitting full use of available light (Maire et al., 2012).

In this work, we propose an extension to the least-cost theory, shifting from the phenomenological β(θ) formulation of soil moisture limitation to a description in which soil water availability increases the carbon cost of water transport (the *a* term as defined by Prentice et al., 2014) and thereby reduces stomatal conductance and χ. First, we define the theoretical value of *a* necessary to sustain carbon assimilation as a function of water use efficiency (WUE). Then, using half-hourly data from a global network of eddy covariance sites, we calculate WUE and quantify empirically how environmental variables affect the value of *a*. This work builds on that of Zhang et al. (2025), who first demonstrated the close relationship between woody stem respiration rates and water transport (as assumed in the least-cost formulation by Prentice et al., 2014) using a combination of field and laboratory measurements and information from experimental manipulations of growth temperature. The present study goes a step further by translating this empirical stem-respiration–water-transport relationship into an ecosystem-scale cost formulation. In doing so, it tests whether variation in the inferred cost of water transport can explain soil-moisture effects on canopy conductance, transpiration and water-use efficiency across sites and climates.

### Theory

The least-cost hypothesis states that plants minimise the combined costs (per unit assimilation) of maintaining transpiration and carboxylation capacities. It leads to the prediction of an optimum χ (Prentice et al., 2014) by balancing the marginal carbon costs of carboxylation capacity and water transport capacity when:

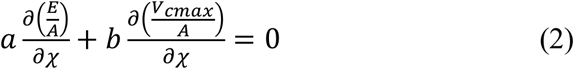

where *a* is the cost of maintaining transpiration to support assimilation at a given rate, conceptualised as the ratio of leaf-specific stem respiration (*R*_s_) to water transport (*S*_W_) (Prentice et al., 2014):

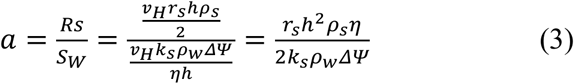

where *v*_*H*_ is the Huber value (ratio of sapwood area to leaf area), *r*_*s*_ is the sapwood-specific respiration rate (*s*^−1^), *h* (m) is the mean length of the path for water transport (roughly equivalent to plant height) , *k*_s_ (*m*^2^) is the sapwood permeability, ρ_w_ and ρ_s_ are water and sapwood molar densities respectively (*mol m*^−3^), *η* (*Pa s*) is the dynamic viscosity of water, and ΔΨ (*Pa*) is the soil-leaf water potential gradient. *b* is the cost of maintaining *V*_cmax_ to sustain assimilation at the same rate, conceptualised as the ratio of leaf dark respiration to V_cmax_. *b* was originally assumed to be constant at 0.015 (Collatz et al., 1991), while *a* was assumed to decline with increasing temperature due to decreasing *η* (Eq. 3). Thus the cost ratio (*b*/*a*) was approximated by β_cost_/η^*^, where β_cost_ = 240 based on leaf δ^13^ C measurements (Wang et al., 2017) – later revised, using a more exact calculation, to 146 (Stocker et al., 2020a) – and η^*^ is the viscosity of water relative to its value at 25°C. *b* can be more accurately approximated as a function of temperature (Ren et al., 2024; Wang et al., 2020a):

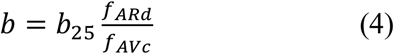

where *b*_25_ = 0.037, constant at any growth temperature (Wang et al., 2020a), represents the ratio of leaf dark respiration at 25 ºC (R_d,25_) to carboxylation capacity at 25 ºC (V_cmax,25_); f_Ard_ is the thermal response function for R_d_ (Heskel et al., 2016); and f_AVc_ is an Arrhenius-type thermal response curve (with deactivation) for V_cmax_ (Kattge and Knorr, 2007).

The least-cost hypothesis predicts the optimum χ as:

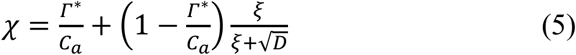

Where

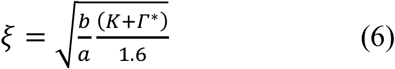

with *K* being the effective Michaelis constant of Rubisco. From Fick’s law, assimilation (*A*) and transpiration (*T*) are given by:

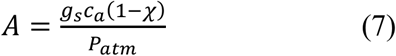

And

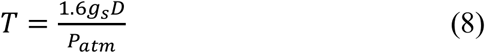

where *P*_atm_ is atmospheric pressure. Equations (7) and (8) lead to:

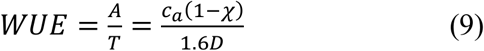

Substituting equations (5) and (6) into equation (9), and solving for the *a*-cost, yields:

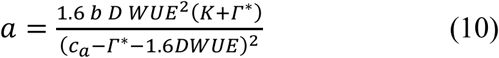

Empirically, WUE can be estimated using eddy-covariance based carbon and water fluxes as:

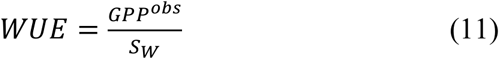

where *GPP*^*obs*^ is GPP at the flux tower, and *S*_*W*_ is the water uptake (assumed to be in steady state with transpiration). The latter was estimated directly from sapflow measurements, if available, or using estimated transpiration to evapotranspiration ratios (*f*_ET_) and latent heat flux (LE) measurements. Actual evapotranspiration (ET) was estimated by converting LE to water flux using the latent heat of vaporisation (*λ*) with corrections for temperature (*T*_air_) and pressure:

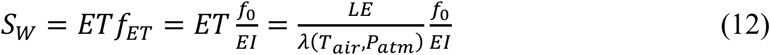

We estimated *f*_*ET*_ as the ratio of *f*_*0*_ to evaporative index (EI), where EI is the ratio of total evapotranspiration to precipitation and *f*_0_ is the fraction of antecedent annual precipitation used as transpiration (*T*/*P*). This quantity shows a hump-shaped response to climatological aridity (Good et al., 2017), which can be predicted using theory developed by Porporato et al. (2004). For simplicity, however, in this work we adopted an empirical relationship (Zhou et al., 2025) derived from the data presented by Good et al. (2017) , using the long-term climatological aridity index (AI = PET/P, where PET is annual potential evapotranspiration and P is annual precipitation) as a predictor:

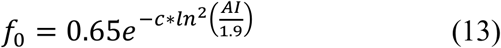

where 0.65 is the maximum *T*/*P*, 1.9 is the AI at which this maximum occurs, and *c* = 0.604169.

The theoretical formulation of S_W_ (equation 3) suggests that its short-term environmental responses (instantaneous to daily) are mostly caused by temperature (via viscosity) and water availability (via the water potential gradient, ΔΨ). Depending on the degree of isohydricity, ΔΨ in general should increase with increasing Ψ_s_ (Martínez-Vilalta et al., 2014), so we expect a strong positive correlation of S_W_ with soil moisture (S_W_’) at this time scale. Responses to long-term bioclimatic variables might therefore involve changes in sapwood permeability more than changes in height, since we expect apex-to-base widening of xylem elements to largely reduce the dependence of water transport on path length (Alemán-Sancheschúlz et al., 2026).

Following the conceptualisation of *a* in equation (3), and using equations (10) and (12), total ecosystem sapwood respiration should be proportional to total ecosystem water transport:

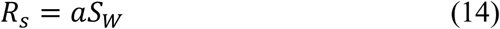

As shown by Zhang et al. (2025) , we expect to see a strong response of *R*s calculated with Eq. 14 to temperature in both, instantaneous and growing season timescales (*R*s’). Then, combining equations (4) and (14), we have:

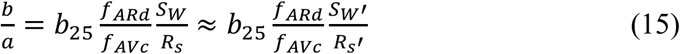

where *S*_W_’ and *R*s’ are the empirical approximations of *S*_W_ and *R*s.

To avoid inconsistencies in the measurements of θ among sites (e.g. different depths of sensors) and the use of any arbitrary thresholds and uncertainties in pedotransfer functions (Paschalis et al., 2022), we used effective soil moisture (Θ^*\**^) derived from flux measurement, calculated as the ratio of ET to PET (Allen, 1998; Davis et al., 2017; Verstraeten et al., 2008), as the index of soil moisture:

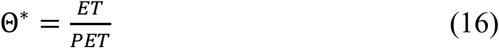

PET was estimated following the Priestley-Taylor model with as modified by Yang & Roderick (2019) to account for surface temperature feedback on net radiation:

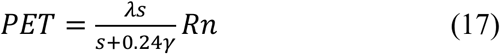

where *s* is the slope of the temperature-saturation vapour pressure curve, *γ* is the psychrometric constant and *R*_n_ is net radiation. Then, replacing equation (17) in (16) and rearranging:

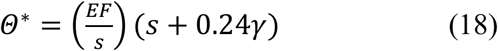

where the evaporative fraction (EF = λ ET/*R*_n_) scales linearly with Θ^*\**^, in agreement with the observations of Fu et al. (2022).

## Materials and Methods

### Global Eddy Covariance and Sapflow data

We used half-hourly observations from global datasets of eddy covariance towers measuring water and carbon fluxes, comprising 709 sites from FLUXNET Shuttle and 105 sites from FluxDataKit, (Hufkens, 2022; Pastorello et al., 2020; Ukkola et al., 2022). Only GPP values derived using the night-time partitioning method were used; gap-filled values were excluded. We added data from those SAPFLUXNET (Poyatos et al., 2020) sites that are collocated with eddy-covariance towers (n = 26). To upscale the sapflow measurements to the ecosystem level, we followed the protocol described by Bright et al. (2022) and Nelson et al. (2020). Savannas, where sapflow measured on sparse trees is not representative of the ecosystem, and sites where the upscaled transpiration exceeded tower-based ET (possibly due to sensor calibration errors) were excluded from the analysis (Fig. S1).

### Global gridded datasets and δ^13^C data

For the global evaluation, monthly forcing data for 2003 to 2020 from ERA5-Land (Muñoz-Sabater, 2019) were used together with fAPAR from the MODIS MCD15A3 product (Myneni et al., 2021) resampled to 0.1°. Effective soil moisture was simulated using SPLASH2 (Sandoval et al., 2024), with soil data from HWSD v2.0 (FAO and IIASA, 2023) used here to estimate soil hydrophysical properties. δ^13^C isotopic data were compiled from open global datasets (Adams et al., 2020; Cornwell et al., 2018; Lavergne et al., 2020a; Mathias and Thomas, 2021; Wang et al., 2026) adding up to 2305 sites (n ∼11k) over the MODIS era. The data were then reprocessed using standard methods implemented in the isocalcR package (Mathias and Hudiburg, 2022), and used to estimate *χ*, adjusting the coefficients according to tissue type (leaf or wood), following Lavergne et al. (2020a). To define the growing season from monthly simulations we used seasonal phase and concentration of GPP following the method described by Kelley et al. (2013).

### Bioclimatic variables

Aridity index was estimated as the ratio:

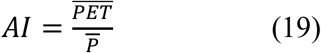

Where 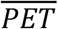 is the mean annual potential evapotranspiration and 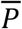 is mean annual precipitation. Growing degree days (GDD_0_) were defined as days when daily mean air temperature >0°C; growth temperature (*T*_G_) was estimated as the average temperature during GDD_0_. All the environmental data used to estimate the bioclimatic metrics were measured *in situ*, except for sites where the records were shorter than five years. In such cases, data from ERA5-Land were used instead.

### Statistical analysis

To obtain a first approximation of the environmental responses of *a* and their functional forms, we evaluated the responses of its two components, *R*_s_ and S_W_ (Eqs. 3, 14), separately using a two-stage approach. In the first stage, mixed-effects models were fitted using the lme4 R package (Bates et al. 2015), using log-transformed dependent variables and site as a random effect. For Rs, instantaneous air temperature was used as a fixed effect, given its known control over respiration rates. For S_W_, we used the ratio of effective soil moisture to relative viscosity 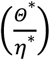, hereafter referred to as “soil-water fluidity”, which captures the combined effect of water availability (via Δψ) and temperature (via viscosity), as predicted by Eq. 3. The fixed effects therefore represent short-term environmental responses, whereas the site-specific coefficients capture long-term adapted responses. In the second stage, the site-specific coefficients were regressed against growth temperature (T_G_) for Rs and aridity index (AI) for S_W_.

A subsequent global calibration of the coefficients found in the previous section was then performed using the full P-model structure and *χ* derived from the δ^13^C data to account for assumptions in the T/ET partitioning and to correct for the unit cost of maintaining J_max_ (c*=0.41), which in the original P-model was estimated assuming *χ* = 0.8 and a constant J_max_:V_cmax_ ratio of 1.88 (Stocker et al., 2020b). The global calibration was performed using the Differential Evolution algorithm (Ardia et al., 2011) with the mean squared error of predictions (MSEP) (Haefner, 2005) as objective function.

## Results

Inferred *R*_s_ and *a* from eddy-covariance data alone show variation over orders of magnitude within and between biomes with no clear overall patterns. However, in general, forests show higher *R*_s_ and *a*, followed by grasslands (GRA) and closed and open shrublands (CHR, OSH). Evergreen broadleaf forests (EBF) were significantly different from evergreen needleleaf forests (ENF) and deciduous broadleaf forests (DBF). Woody savannas, mostly occurring at sites with high seasonality, show the highest *R*_s_ and *a*, whereas Wetlands (WET) and Polar tundra (SNO) show the lowest *R*_s_ and *a* (Figs. 1B, 1D).

**Fig. 1.**
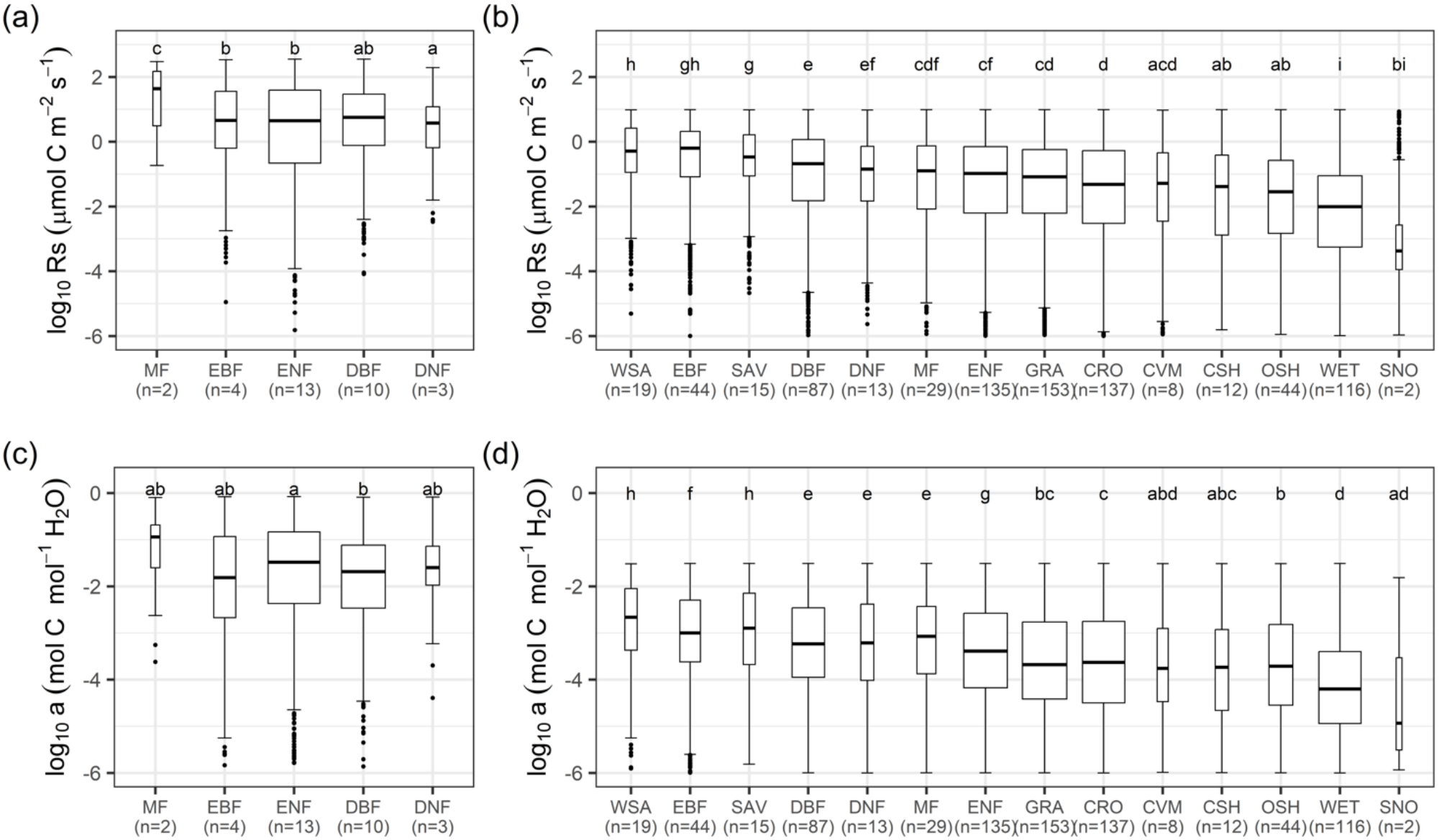
Carbon costs of water transport in different biomes. Boxes show the interquartile range with the median indicated by the thick solid line. Whiskers extend to the most extreme values within 1.5× the interquartile range, and outliers are shown as black circles. Box widths are proportional to sample size. Post-hoc Tukey test results are shown at the top of the bars. **A** ecosystem-sapwood respiration from sap flow and eddy covariance. **B** ecosystem-specific sapwood respiration from eddy covariance alone. **C** Carbon cost of transpiration estimated from sap flow and eddy covariance. **D** Carbon cost of transpiration estimated from eddy covariance alone. ENF stands for Evergreen Needleleaf Forests, EBF for Evergreen Broadleaf Forests, DNF for Deciduous Needleleaf Forests, DBF for Deciduous Broadleaf Forests, MF for Mixed Forests, CSH for Closed Shrublands, OSH for Open Shrublands, WSA for Woody Savannas, SAV for Savannas, GRA for Grasslands, WET for Permanent Wetlands, CRO for Croplands, CVM for Cropland/Natural Vegetation Mosaics, and SNO for lands under snow/ice cover most of the year.

Sapflow estimates of transpiration (Figs 1A and 1C) show that, in general, *Rs* and *a* are slightly higher in evergreen forests than deciduous. Moreover, both quantities are highest in Mixed forests (MF) and smallest in Deciduous needleleaf forests (DNF). However, the small number of sites makes interpretation difficult.

Inferred *R*_s_ from EC data was strongly dependent on temperature, as expected. The short-term response (Eq. 20a), increased with instantaneous temperature (marginal R^2^ = 0.52, *p* < 0.001) (Fig. 2A) – opposite to the long-term response (Eq. 20b), which decreased with growth temperature (R^2^ = 0.05, *p* < 0.001) (Fig. 2B). DNF and ENF sites show the highest long-term *R*_s_; while WET and CVM showed the lowest.

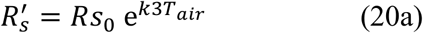

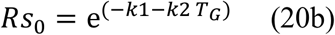

**Fig. 2.**
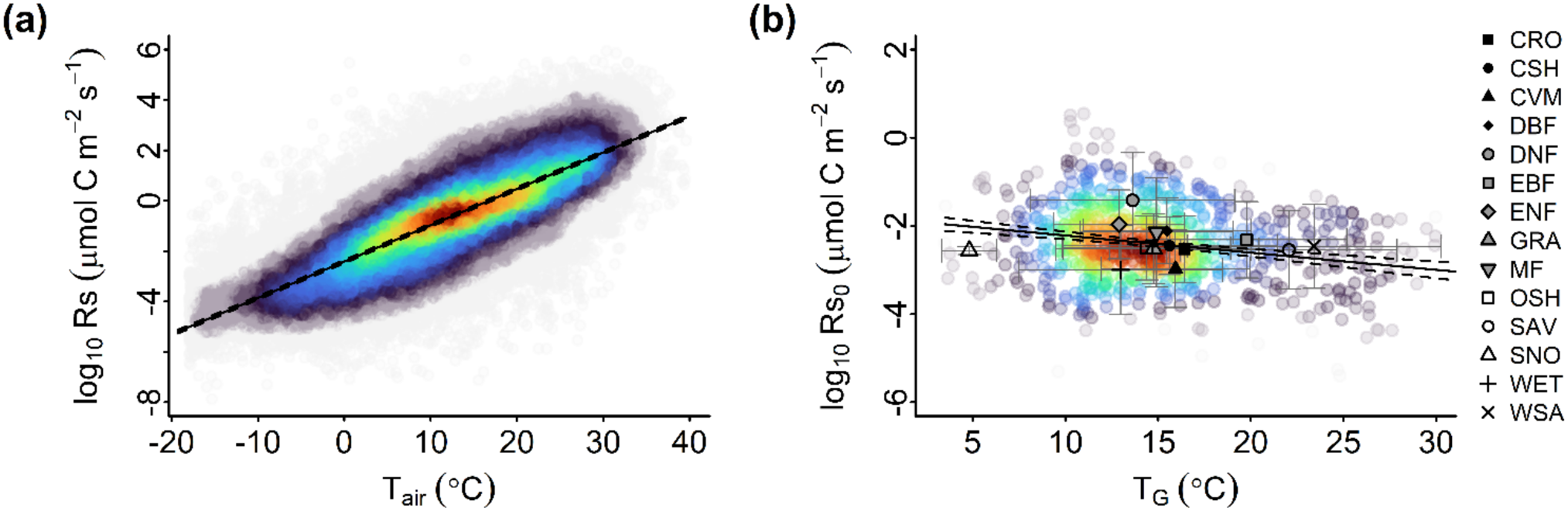
Inferred ecosystem-level sapwood respiration response to temperature derived from EC data. **A.**Instantaneous response to air temperature (partial residual plot). **B**. Long-term response to growth temperature.

In the short term, water transport shows a strong proportional increase with soil-water fluidity based on within-site variation (marginal R^2^ = 0.53, *p* < 0.001) (Eq. 21a, Fig. 3A):

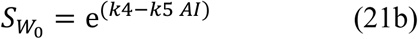

Whereas in the long-term, water transport shows only a weak response to increasing (R^2^ = 0.21,

**Fig. 3.**
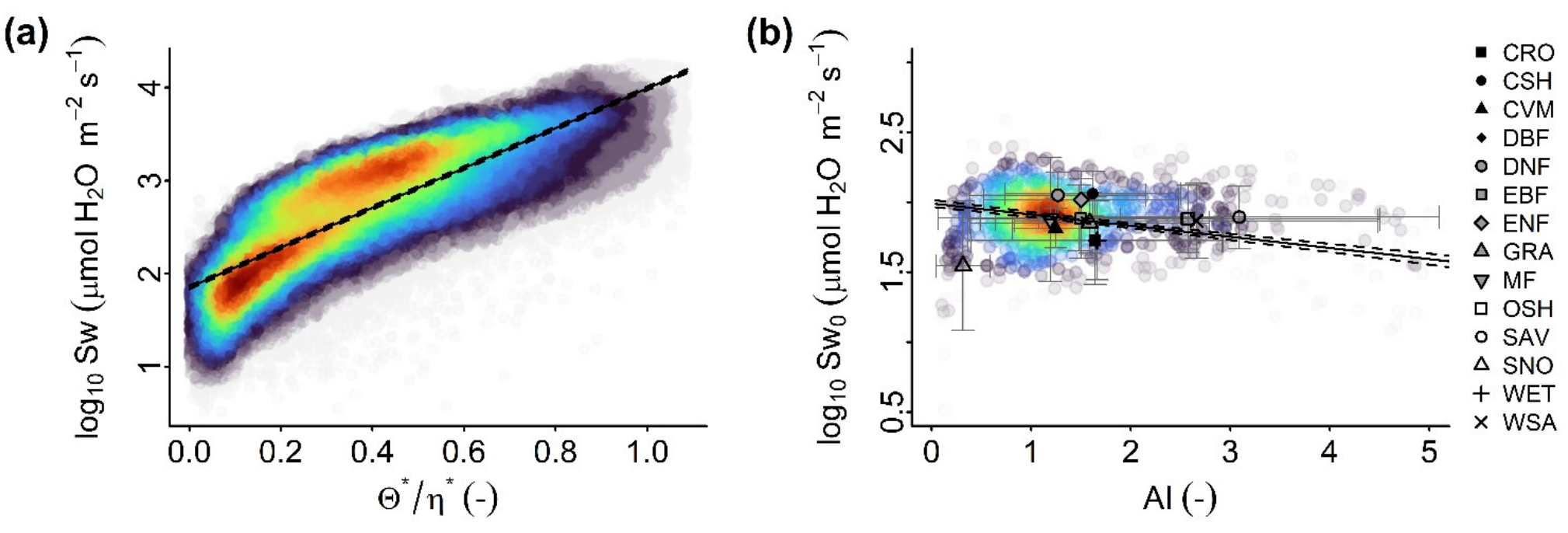
Water transport response to water availability derived from EC data. **A** Instantaneous response to soil-water fluidity (partial residual plot). **B** Long-term response to aridity index (AI).

*p* < 0.001):

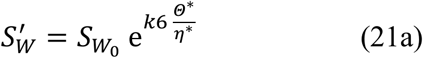

Polar tundra sites (SNO) show the lowest long-term *S*_w_, whereas CSH displays the highest. SAV, WSA and CSH show similar values of long-term *S*_w_ while spanning most of the aridity spectrum (Fig. 3B).

Optimizing the new formulation using *χ* derived from δ^13^C data (Table 1) improved the slope of the observed vs. simulated regression, from 0.31 to 0.39 compared with the original P-model formulation, along with a substantial increase in the R^2^ from 0.09 to 0.16 (Figs. 4B, 4C). This improvement reflects a correction of the overestimation of *χ* during the growing season in grasslands in Africa and Asia, and in open shrublands in the Americas and Australia, while in tropical EBF and boreal CSH the new formulation corrected the previous underestimation (Fig. 4A).

**Table 1.** Coefficients of the empirical approximations of R_s_ and S_W_ after the global optimization.

| k1 | k2 | k3 | k4 | k5 | k6 |
|---|---|---|---|---|---|
| 2.559 | 0.099 | 0.080 | 3.99 | 0.005 | 2.003 |

**Fig. 4.**
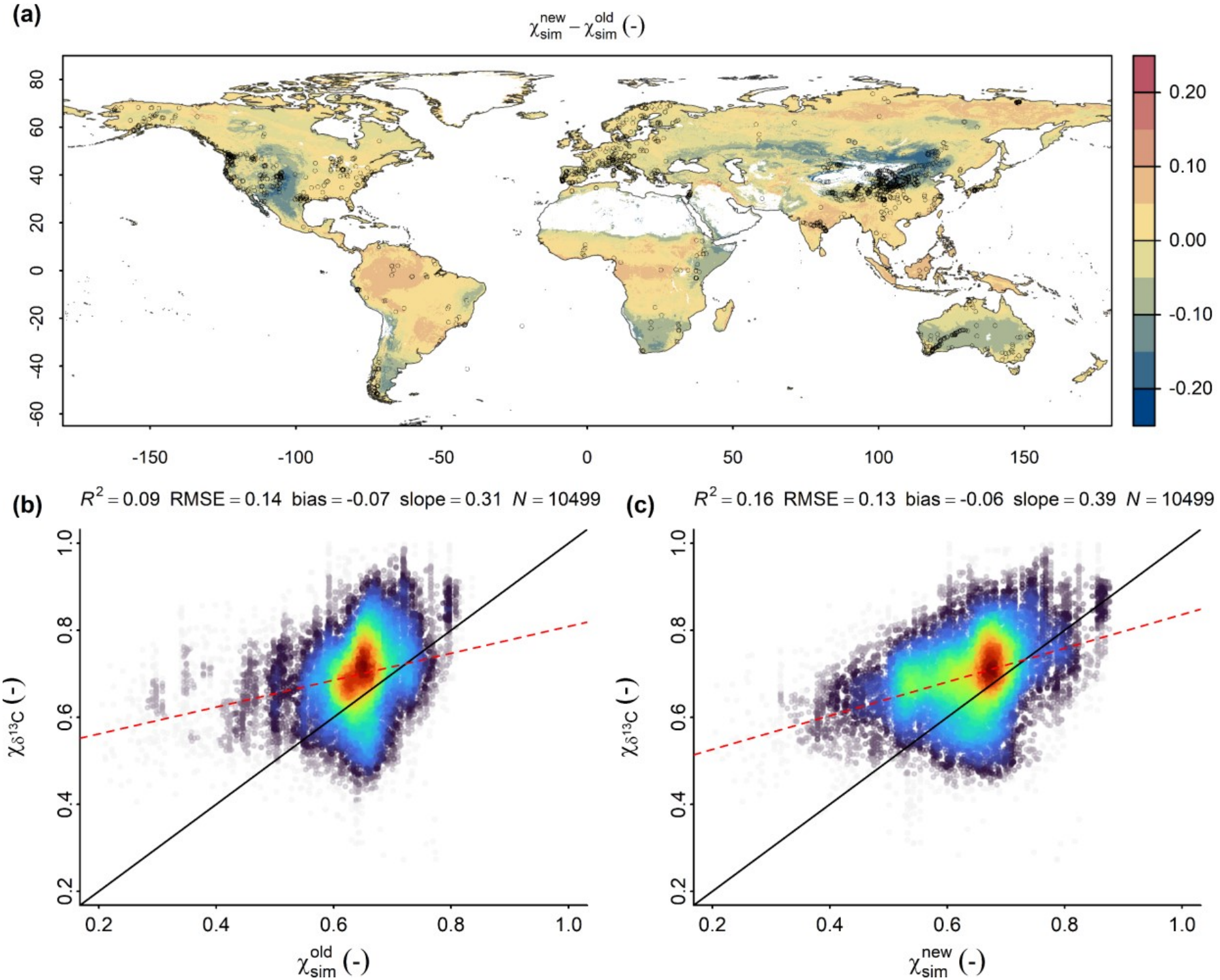
Global performance of the P-model with the new b/a cost schema simulating *χ*. **A** Spatial distribution of the difference between new and the original P-model simulations of growing season *χ*, averaged over the years 2003 to 2020. Small circles show the sampling sites of δ^13^C **B** Correlations of δ^13^C based *χ* and simulated values using the original formulation. **B** Correlations of δ^13^C based *χ* and simulated values using the new formulation.

When canopy conductance (*G*_c_) derived from the sapflow observations, by rearranging Eq. 8, was compared to the predictions of the least-cost hypothesis using the original formulation 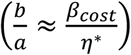, we found that the original form underestimated low values of *G*_c_ and overestimated high values of *G*_c_ (Fig 5A). The new conceptualization of *b*/*a* using equation (15) increased the *R*^2^ for canopy conductance predictions from 0.34 to 0.52 and increased the slope of the observed versus simulated regression from 0.29 to 0.52 (Fig 5B). The prediction of transpiration also improved, with *R*^2^ increasing from 0.52 to 0.65 and the slope of the observed versus simulated regression increasing from 0.35 to 0.57 (Fig 5D).

**Fig. 5.**
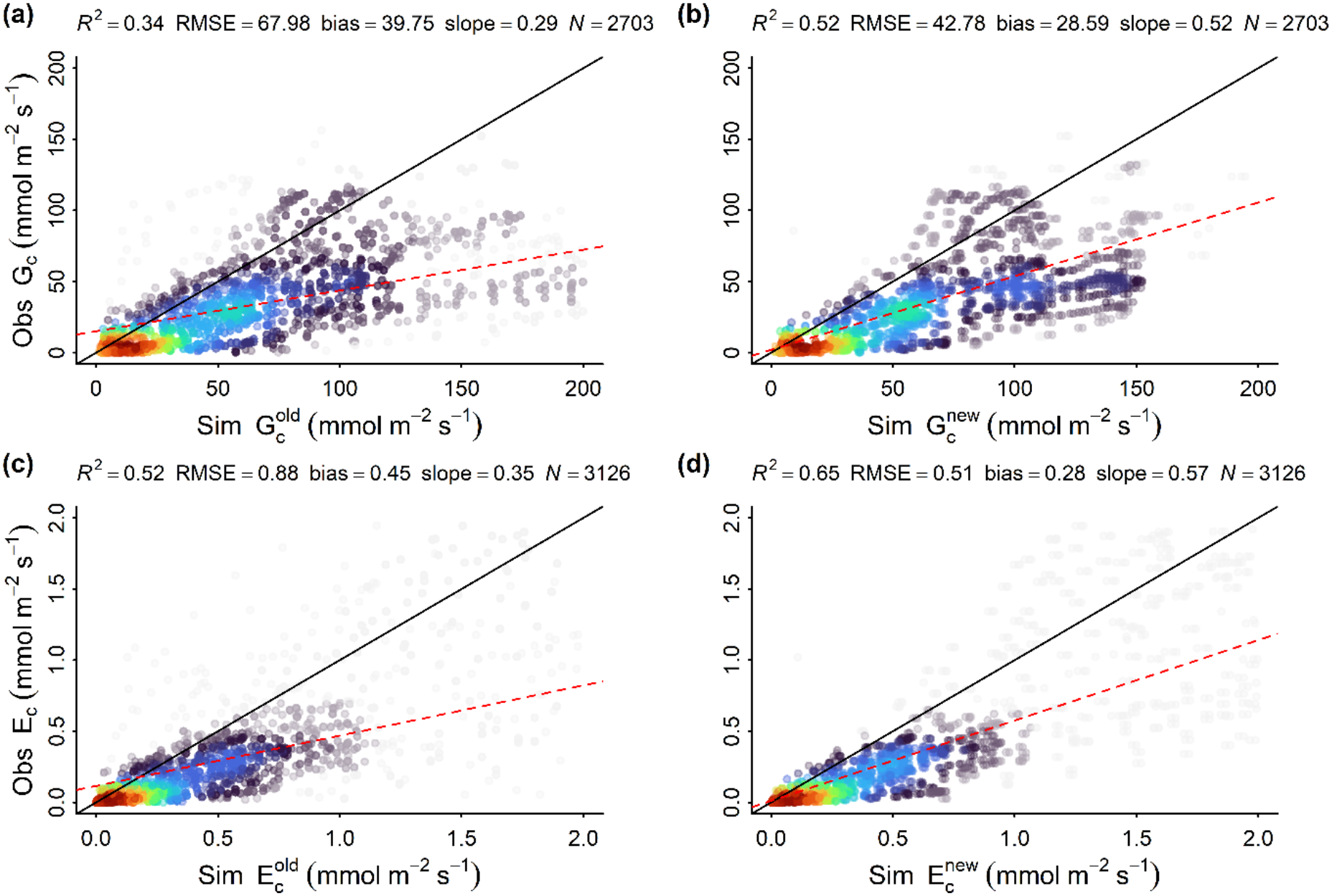
Correlations of sap flow-based observed and simulated values. **A** Canopy conductance simulated with the original formulation of costs. **B** Canopy conductance simulated using equation (15). **C** Transpiration simulated with the original formulation of costs. **D** Transpiration simulated using equation (15).

Although the statistics in Figures 5C and 5D generally show better agreement with observations than the original model, a more detailed site-by-site analysis shows that, while the shape of the seasonal cycle is captured in all sites, both formulations slightly overestimate the magnitude in almost every site but underestimate transpiration at FR-Pue and IL-Yat (Fig. 6).

**Fig. 6.**
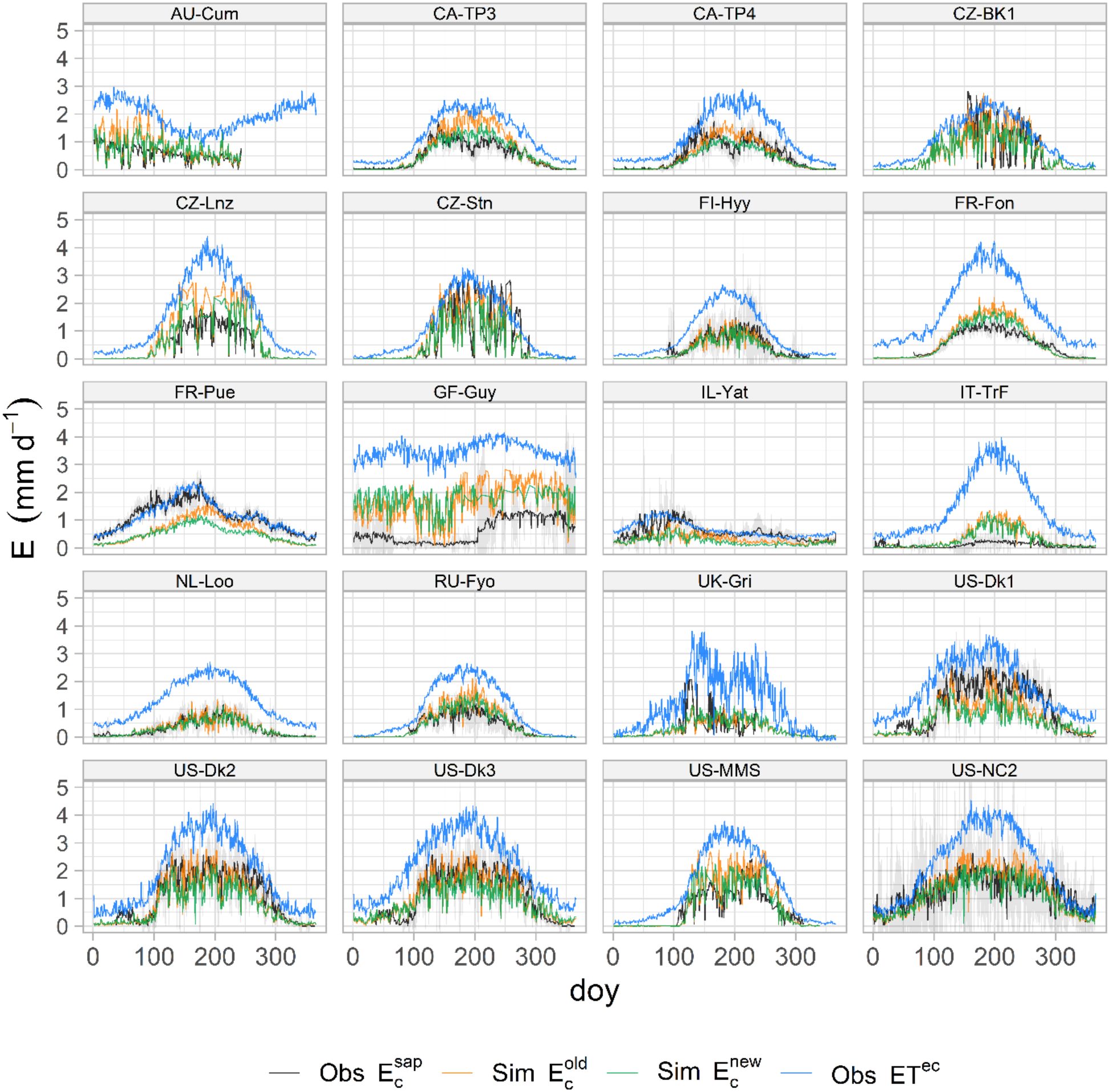
Observed and simulated transpiration at selected SAPFLUXNET sites. The black continuous line shows the mean observed transpiration along the day of the year (doy), with grey ribbons representing the standard deviations of the observations. The green line shows the simulated transpiration using equation (15); the orange line shows the simulated transpiration using the original formulation of the costs (Stocker et al. 2020) and the blue line evapotranspiration from the tower.

## Discussion

In this work, we propose an extension to the least-cost theory by incorporating soil moisture control on water and carbon fluxes via the carbon cost of water transport (*a*), thereby influencing canopy conductance (*G*_c_). We posit that low soil moisture increases *a*, which in turn reduces canopy conductance, thereby lowering both assimilation and transpiration. We tested this idea against independent sapflow measurements and found that it substantially increases the explanatory power of the least-cost theory in predicting *G*_c_ and transpiration.

However, our results also suggest that the general overestimation by both versions is most likely due to missing processes that modify the cost ratio *b/a*. The value of *b*_25_ reported by Wang et al. (2020) does not vary systematically with climate but shows substantial unexplained variation, supporting this interpretation.

On the one hand, results by Paillassa et al. (2020) suggest that soil pH modifies *b/a*. Under acidic conditions, the solubilization of toxic manganese and aluminum increases the respiratory costs associated with detoxification. Conversely, under slightly alkaline conditions, greater magnesium availability may increase photosynthetic biochemical capacity by reducing the costs of acquiring nitrogen and other nutrients. Cheaib et al. (2025) have further proposed that variations in *b/a* are driven by nitrogen availability, via mycorrhizal associations.

On the other hand, Walker et al. (2014) showed that leaf phosphorus (P) substantially increases the sensitivity of *V*_cmax_ to leaf nitrogen (N). Maire et al. (2015) showed that species growing on high-P soils tend to increase their maximum photosynthetic rates to compensate for lower stomatal conductance. Both studies therefore suggest that *b/a* is influenced by P availability.

The spatial pattern of the difference between χ predicted using the original and new schemes appears similar to the difference between χ predicted using climate-only and a climate-soil interactive model from Paillassa et al. (2020). This suggests that the water holding capacity derived from SPLASH2 is capturing some of the variation attributable to interactions between soil texture and water supply.

Although the overall patterns are supported by independent sap-flow based evaluation, we note that the inference of WUE, *a* and ecosystem-level Rs depends partly on the partitioning of evapotranspiration into transpiration. For sites without direct sap-flow observations, this partitioning relies on an empirical *f*_0_ relationship, which might introduce uncertainty in the magnitude of inferred water transport costs.

We also found that the inferred short-term response of sapwood respiration at ecosystem scale increases with increasing temperature, while the long-term response (Rso) decreases with growth temperature. These patterns are fully consistent with measurements of stem respiration in woody plants (Zhang et al., 2025).

Soil-water fluidity explained most of the variation in water transport. This finding is consistent with hydraulic theory, via Darcy’s law, which establishes this relationship to be proportional to the ratio ΔΨ/η (equation (3)). *S*_w_ varied across several orders of magnitude, consistent with the known variability on sapwood conductivity (Sperry, 2003). Nevertheless, this response conveys the idea that it is not just the amount of water available that matters, but also how easily that water can be transported — a function of viscosity and xylem vessel properties — which could explain higher sensitivities to soil moisture in arid and warm sites, as observed by Giardina et al. (2023).

The ratio *R*_s_/*S*_W_ then would theoretically reflect maintenance respiration for all tissues involved in water transport, potentially encompassing both the costs of short-term active regulation and long-term passive structural support. Thus, in the short-term, this ratio may reflect the ATP used to produce organic solutes (Sperry, 2003) aiming to maintain turgor pressure (or leaves RWC) via active ion transport (e.g., K^+^ pumps) across membranes. This energy-expensive active control can also increase xylem pressure to help reverse cavitation and rehydrate leaves after drought stress (Venturas et al., 2017). However, since most of the water-loss regulation occurs in the leaf at a sub-daily time scale and that embolism is uncommon and may be limited to periods of severe drought (Lens et al., 2022), it may be that a larger portion of the cost of maintaining water transport capacity is spent on sustaining turgor, rather than restoring xylem conductivity. Since most models assume a steady state, they cannot reproduce this process (Buckley, 2019). Another potential cost could be the production of antioxidant enzymes, including superoxide dismutase, catalase and peroxidase, to neutralize reactive oxygen species produced during dehydration.

Our estimated values of *a* were consistently higher in woody than herbaceous species (Fig 1), possibly reflecting the long-term cost of sustaining tougher cell walls and wider and longer xylem vessels (Olson et al., 2018). This difference might also reflect seasonal root shifts to deeper soil layers – a strategy observed globally that allows plants to maintain transpiration and hydraulic safety under drought conditions (Bachofen et al., 2024b). It could also include the increase of aquaporin activity in roots and leaves to enhance water uptake.

We observed a weak but significant increase of *a* with aridity (through decreasing Sw), suggesting that water transport is more expensive in arid regions than in wet regions. However, our results also show *T*_G_ to have a stronger effect than AI (see equation (21b)), suggesting that it is more expensive to transport water in colder climates than in arid regions. Thus, in cold climates, shorter plants with narrow vessels that are less costly to maintain would have an advantage (Olson et al., 2018). This contrasts with the suggestion by Prentice et al. (2014), that plants may adapt to arid regions by lowering their height and thus lowering *a*. Almost unchanged sapwood-specific hydraulic conductivity (*K*s) with increasing aridity (Zhu and Zhao, 2023) also support this idea.

The weak effect of AI on *a* and (hence on *G*_c_) further suggests that the strong reduction in maximum light use efficiency (LUE) observed in arid sites (Bassiouni et al., 2023; Mengoli et al., 2025; Sandoval et al., 2026b) is not primarily due to *G*_c_, but mostly due to non-stomatal responses. Consistent with this idea, Bassiouni et al. (2023) proposed that the synchronisation between water uptake and transpiration increases with increasing aridity. Since this synchronisation is mediated by higher leaf water capacitance, larger elastic moduli, or lower osmotic potentials – all of which are costly to produce – the carbon investment in photosynthetic proteins in the long-term is likely to be reduced while the cost in hydraulic structures is amortised.

Finally, our new revised formulation of the ratio *b*/*a* implies three key predictions in response to decreasing soil water availability, with all else equal. The first is an increase in iWUE, this is consistent with observations from Driscoll et al. (2020); Li et al. (2022); Jiang et al. (2024) and Wang et al. (2026) (Fig. 7a). The second is a decrease of *ξ* (analogue to g1 in the Medlyn et al. (2011) model), this has also been observed experimentally before a drop in *V*_*cmax*_ at extremely dry soil conditions (Zhou et al., 2013, 2014) (Fig. 7b). The third prediction is a slight increase in *V*_*cmax*_, as shown for species adapted to a dry climate when subjected to moderate drought (Zhou et al., 2016), followed by a decrease as Θ^*\**^ approaches zero. The latter appears to be the case for most of the species analysed by Zhou *et al*. (2013, 2014). Although in our model this decrease emerges when D is high (Fig. 7c), our scheme is still unable to produce predictions beyond the lower limit of the effective soil moisture, where the dramatic drop in *V*_*cmax*_ occurs (Ψ_pd_ < -1.5 MPa) (Zhou et al., 2013, 2014).

**Fig. 7.**
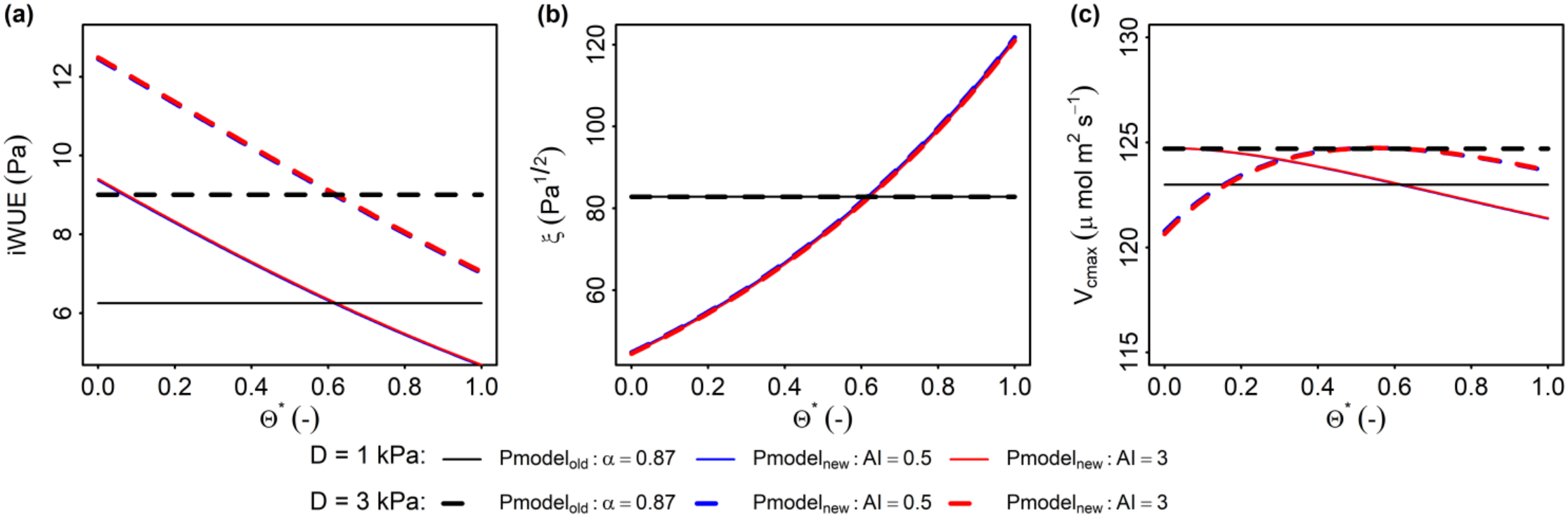
Predictions of the new b/a scheme. **(a)** iWUE at different soil moisture and aridity levels. **(b)** ξ at different soil moisture and aridity levels. **(c)** *V*_*cmax*_ at different soil moisture and aridity levels. Simulations were run at sea level, a temperature of 25 ºC, CO_2_ = 400 ppm and PPFD = 1500 μmol m^−2^ s^−1^ with D at 1kPa and 3kPa.

Overall, our idea that low soil moisture increases *a*, which in turn reduces canopy conductance and thereby lowers both assimilation and transpiration, significantly improves the predictive power of the least-cost hypothesis, offering a more mechanistically coherent alternative to existing parameterisations of soil moisture limitation, that are crucial for predicting water and carbon fluxes future scenarios.

## Code and data availability

The modified P-model code with the new formulations proposed here is available on Zenodo at https://zenodo.org/records/24084280 (Sandoval et al., 2026a). The code to reproduce the results presented here is available at https://zenodo.org/records/24084318 (Sandoval, 2026). Eddy covariance data used in this study is openly available at https://data.fluxnet.org/data/ and FluxDataKit at https://doi.org/10.5281/zenodo.12818273.

## Author contributions

DS: Conceptualisation, data analysis and first draft writing. VF: scientific input, code supervision, HZ: scientific input, ICP: scientific input, supervision. All the authors reviewed and edited the final manuscript.

## Competing interests

None declared

## Acknowledgments

This work is a contribution to the LEMONTREE (Land Ecosystem Models based On New Theory, obseRvations and ExperimEnts) project.

## Financial support

This research received support through Schmidt Sciences, LLC (Grant number G-21-61881: DSC, ICP, HZ).

## Notes

### Competing Interest Statement

The authors have declared no competing interest.

### Summary of Updates

Code and data availability: The modified P-model code with the new formulations proposed here is available on Zenodo at https://zenodo.org/records/24084280 (Sandoval et al., 2026a). The code to reproduce the results presented here is available at https://zenodo.org/records/24084318 (Sandoval, 2026). Eddy covariance data used in this study is openly available at https://data.fluxnet.org/data/ and FluxDataKit at https://doi.org/10.5281/zenodo.12818273.

