## Supplementary Material for "Extending the least-cost theory of stomatal regulation to include soil moisture stress"

Sandoval, D. *et al.*

**This PDF file includes:**

Table S1.
Figs. S1

**Table S1. SAPFLUXNET sites collocated with EC sites.** SPF_id is the SAPFLUXNET ID code, EC_id the FLUXNET/FluxDatakit ID code, IGBP the biome abbreviation, Lon. is Longitude (°) and Lat is latitude (°).

| SPF_id | Name | EC_id | IGBP | Lon | Lat |
|---|---|---|---|---|---|
| AUS_RIC_EUC_ELE | Richmond NSW EucFACE | AU-Cum | EBF | 150.7403 | -33.6178 |
| AUS_WOM | WombatStateForest | AU-Wom | EBF | 144.0944 | -37.4222 |
| CAN_TUR_P39_POS | TUR | CA-TP3 | ENF | -80.3574 | 42.70978 |
| CAN_TUR_P39_PRE | TUR | CA-TP3 | ENF | -80.3574 | 42.70978 |
| CAN_TUR_P74 | TUR | CA-TP3 | ENF | -80.3483 | 42.70681 |
| CAN_TUR_P39_POS | TUR | CA-TP4 | ENF | -80.3574 | 42.70978 |
| CAN_TUR_P39_PRE | TUR | CA-TP4 | ENF | -80.3574 | 42.70978 |
| CAN_TUR_P74 | TUR | CA-TP4 | ENF | -80.3483 | 42.70681 |
| CZE_BIK | Bik | CZ-BK1 | ENF | 18.53 | 49.49444 |
| CZE_LAN | Lanžhot | CZ-Lnz | MF | 16.94639 | 48.68167 |
| CZE_RAJ_RAJ | Rajec | CZ-RAJ | ENF | 16.70129 | 49.44153 |
| CZE_STI | Stitna nad Vlari | CZ-Stn | DBF | 17.97 | 49.03583 |
| FIN_HYY_SME | Hyytiala Forest Field Station | FI-Hyy | ENF | 24.29477 | 61.84742 |
| FRA_FON | Fontainebleau-Barbeau | FR-Fon | DBF | 2.78014 | 48.47634 |
| FRA_HES_HE1_NON | Hesse | FR-Hes | DNF | 7.0647 | 48.6742 |
| FRA_HES_HE2_NON | Hesse | FR-Hes | DNF | 7.0647 | 48.6742 |
| FRA_PUE | Puechabon | FR-Pue | EBF | 3.595833 | 43.74139 |
| GUF_GUY_GUY | Guyaflux | GF-Guy | EBF | -52.9249 | 5.278772 |
| GUF_GUY_ST2 | Guyaflux | GF-Guy | EBF | -52.9122 | 5.281667 |
| ISR_YAT_YAT | Yatir | IL-Yat | ENF | 35.0515 | 31.345 |
| ITA_TOR | Torgnon | IT-TrF | DNF | 7.56089 | 45.82376 |
| NLD_LOO | Loobos | NL-Loo | ENF | 5.743553 | 52.16648 |
| RUS_FYO | Fyodorovskoye | RU-Fyo | ENF | 32.92208 | 56.46153 |
| GBR_ABE_PLO | Aberfeldy | UK-Gri | ENF | -3.8 | 56.616 |
| USA_DUK_HAR | Duke Blackwood Hardwood | US-Dk1 | DBF | -79.1004 | 35.9736 |
| USA_DUK_HAR | Duke Blackwood Hardwood | US-Dk2 | DBF | -79.1004 | 35.9736 |
| USA_DUK_HAR | Duke Blackwood Hardwood | US-Dk3 | DBF | -79.1004 | 35.9736 |
| USA_INM | INMMSF | US-MMS | DBF | -86.4132 | 39.32323 |
| USA_MOR_SF | Morgan-Monroe State Forest | US-MMS | DBF | -86.4133 | 39.32301 |
| USA_PAR_FER | Parker Tract | US-NC2 | ENF | -76.6679 | 35.80311 |
| USA_CHE_ASP | ChEAS | US-PFn | MF | -90.267 | 45.937 |
| USA_CHE_MAP | ChEAS | US-PFn | DBF | -90.26 | 45.948 |
| USA_WIL_WC1 | Willow Creek | US-WCr | DBF | -90.0867 | 45.81306 |
| USA_WIL_WC2 | Willow Creek | US-WCr | DBF | -90.0867 | 45.81306 |

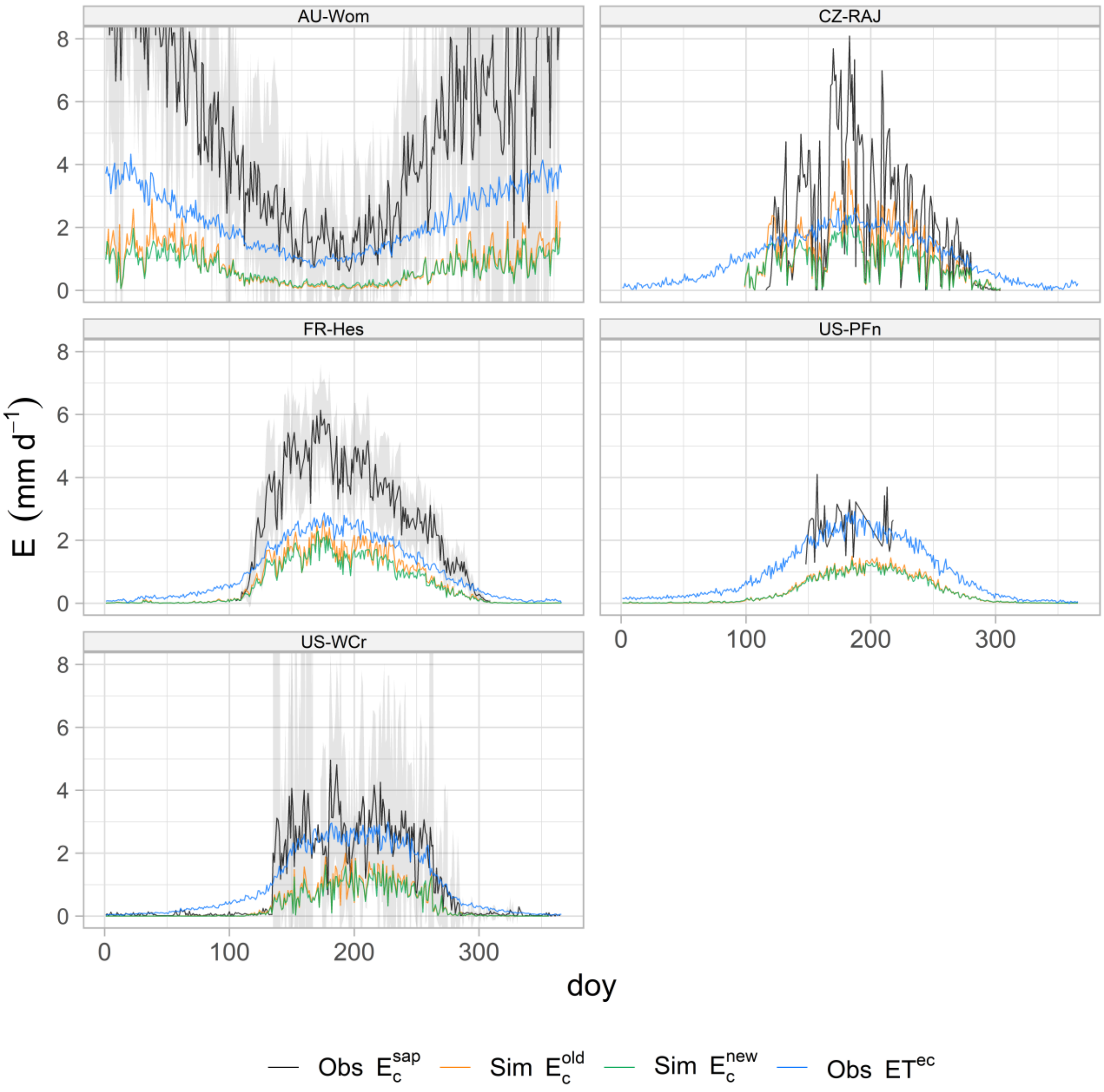


**Figure. S1. Excluded SAPFLUXNET sites.** The black continuous line shows the mean observed transpiration along the day of the year (doy), with grey ribbons representing the standard deviations of the observations. The green line shows the simulated transpiration using equation (15); the orange line shows the simulated transpiration using the original formulation of the costs (Stocker et al. 2020) and the blue line evapotranspiration from the tower.
